# Transcriptomic characterization of human lateral septum neurons reveals conserved and divergent marker genes across species

**DOI:** 10.1101/2024.04.22.590602

**Authors:** Robert A. Phillips, Seyun Oh, Svitlana V. Bach, Yufeng Du, Ryan A. Miller, Joel E. Kleinman, Thomas M. Hyde, Stephanie C. Hicks, Stephanie C. Page, Keri Martinowich

## Abstract

The lateral septum (LS) is a midline, subcortical structure, which regulates social behaviors that are frequently impaired in neurodevelopmental disorders including schizophrenia and autism spectrum disorder. Mouse studies have identified neuronal populations within the LS that express a variety of molecular markers, including vasopressin receptor, oxytocin receptor, and corticotropin releasing hormone receptor, which control specific facets of social behavior. Despite its critical role in regulating social behavior and notable gene expression patterns, comprehensive molecular profiling of the human LS has not been performed. Here, we conducted single nucleus RNA-sequencing (snRNA-seq) to generate the first transcriptomic profiles of the human LS using postmortem human brain tissue samples from 3 neurotypical donors. Our analysis identified 5 transcriptionally distinct neuronal cell types within the human LS that are enriched for *TRPC4*, the gene encoding Trp-related protein 4. Differential expression analysis revealed a distinct LS neuronal cell type that is enriched for *OPRM1*, the gene encoding the µ-opioid receptor. Leveraging recently generated mouse LS snRNA-seq datasets, we conducted a cross-species analysis. Our results demonstrate that *TRPC4* enrichment in the LS is highly conserved between human and mouse, while *FREM2*, which encodes FRAS1 related extracellular matrix protein 2, is enriched only in the human LS. Together, these results highlight transcriptional heterogeneity of the human LS, and identify robust marker genes for the human LS.

## INTRODUCTION

The lateral septum (LS) is a subcortical brain region located along the medial boundaries of the lateral ventricle that controls several aspects of social behavior. Human fMRI studies and clinical case reports demonstrated LS involvement in social cognitive processes such as moral sentiments [1–3], social attachment [4], and sexual behaviors [5], while functional studies in rodents demonstrated that molecularly-defined subpopulations of LS neurons critically regulate social behaviors [6–11] and mediate stress-induced social deficits [12]. To more comprehensively characterize the molecular diversity of these LS cell types, recent single nucleus RNA-sequencing (snRNA-seq) studies in mice identified dozens of unique neuronal subtypes that are both transcriptionally and spatially distinct [13–16]. While snRNA-seq studies of the whole human brain have identified some septal cells [17], no studies to date have specifically targeted the human LS for molecular and transcriptional profiling. Understanding the molecular composition of the LS at single nucleus resolution is essential for understanding the neural substrates that control human social behaviors, many of which are impaired in neurodevelopmental and neuropsychiatric disorders, including autism spectrum disorder (ASD) [18–20], schizophrenia [21], and bipolar disorder (BPD) [22,23].

To understand the molecular diversity of the human LS, we generated snRNA-seq data from postmortem human LS tissue (*N*=3 brain donors). We identified 25 transcriptionally distinct cell clusters with 5 neuronal subtypes originating from the LS. These neuronal clusters exhibited high expression of transient receptor potential cation channel subfamily C member 4 (*TRPC4)*, which has also been identified as an LS marker in mouse snRNA-seq studies [13], suggesting potential conservation of transcriptional markers across species. Cross-species analysis confirmed *TRPC4* as a transcriptional marker of the LS in humans and mice, and identified FRAS1 related extracellular matrix 2 (*FREM2*) as a human-specific marker. Single molecule *in situ* hybridization (smFISH) studies confirmed *FREM2* expression in the LS of an independent donor. This dataset is a resource for understanding molecular heterogeneity within the human LS and spurring novel avenues of research, particularly in understanding the spatial topography of transcriptionally distinct LS neuronal subtypes. To facilitate further exploration of this dataset, we generated a freely available interactive web-based app located at https://libd.shinyapps.io/LS_snRNAseq/.

## RESOURCE AVAILABILITY

### Lead Contact

Further information and requests for resources not provided below (see data and code availability) should be directed to the lead contacts, Keri Martinowich,Stephanie Cerceo Page, and Stephanie C. Hicks.

### Materials availability

This study did not generate new reagents or materials.

### Data and code availability

- All data are publicly available through the sequence read archive with project accession listed above. In addition to raw sequencing files, we provide a SingleCellExperiment object that contains cellular identities, raw counts, and log-normalized counts.
- The code for this project is publicly available and listed in the key resources table. Analyses were performed with R v4.4.0 and Bioconductor v3.19.
- Any additional information required to reanalyze the data reported in this paper is available from the lead contact upon request.

## MATERIALS AND METHODS

### Key Resources Table

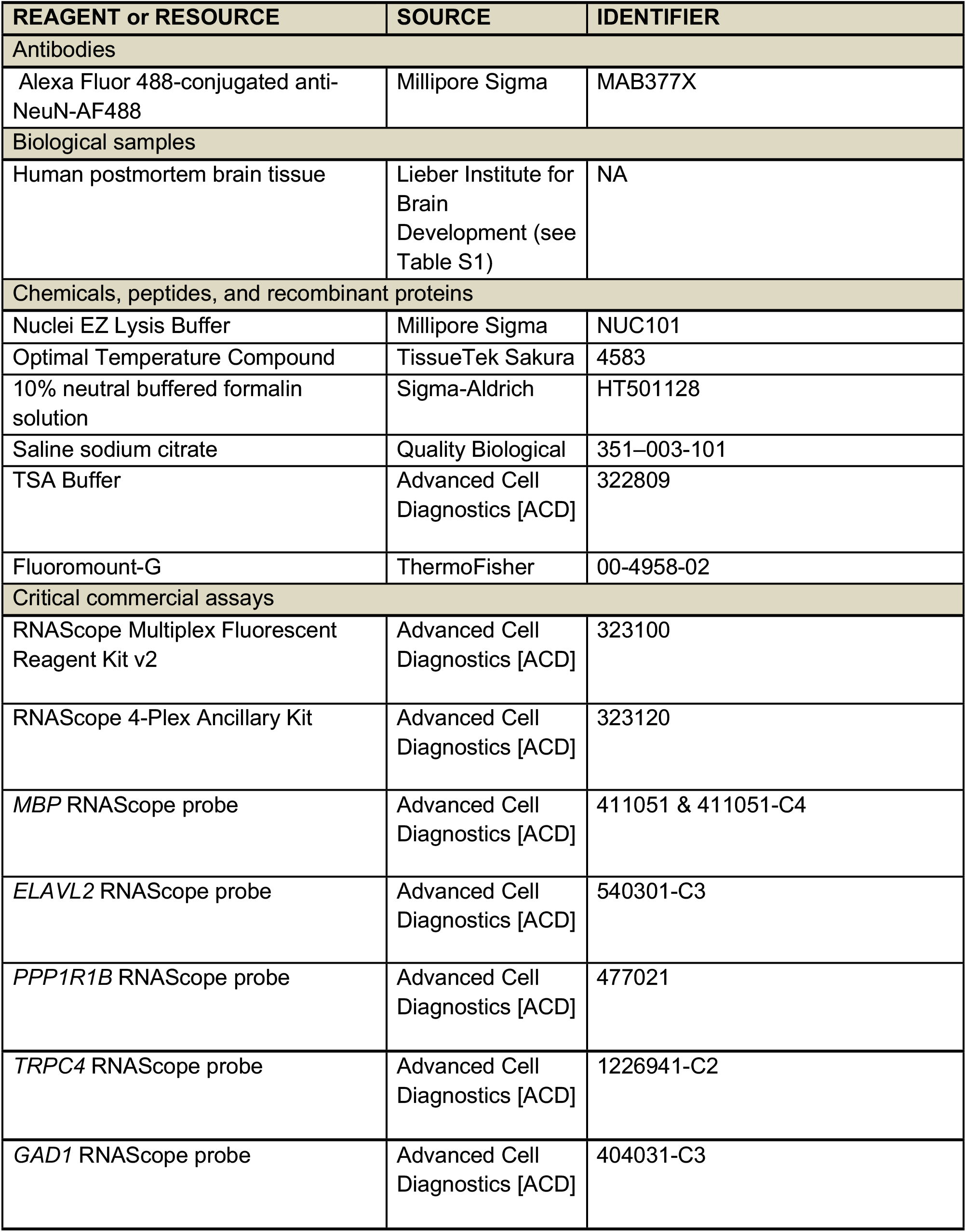
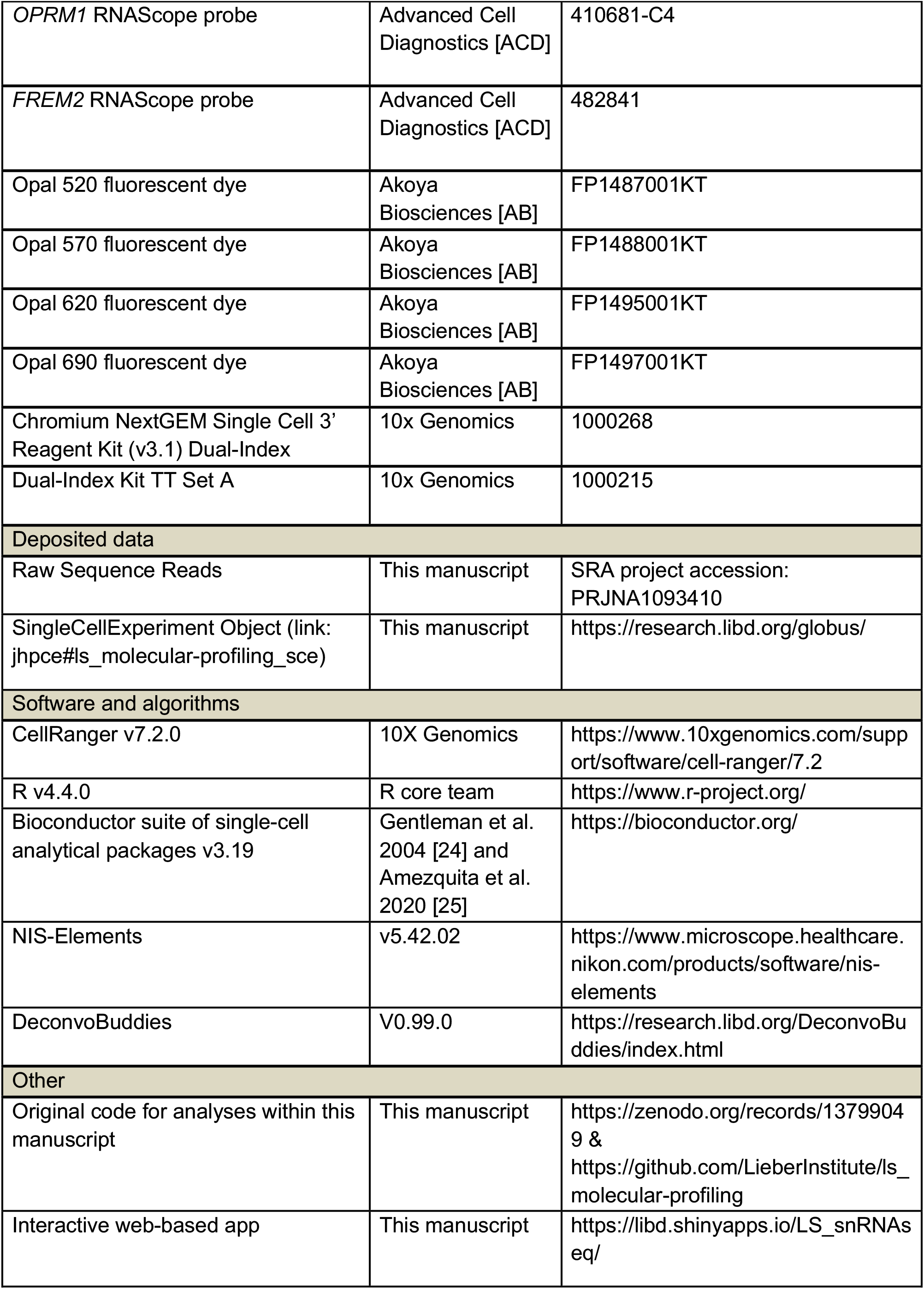

### Postmortem Human Tissue Samples

Postmortem human brain tissue from adult male donors (*N*=4) of Caucasian, African, or Hispanic ancestry spanning ages 47-62 years were obtained at the time of autopsy following informed consent from legal next of kin, through the Maryland Department of Health IRB protocol #12–24 and from the Department of Pathology at Western Michigan University Homer Stryker MD School of Medicine, under the WCG protocol #20111080. Details regarding tissue acquisition, processing, dissection, clinical characterization, diagnosis, neuropathological examination, RNA extraction, and quality control (QC) measures have been previously published [26]. Using a standardized protocol, all donors were subjected to clinical characterization and diagnosis. Macro- and microscopic neuropathological examinations were performed, and subjects with evidence of obvious neuropathology were excluded. All donors were negative for illicit drugs of abuse. Demographics for donors included in the study are listed in **Table S1**. To dissect the LS, fresh frozen coronal brain slabs with clearly visible caudate nucleus, putamen, internal capsule, anterior commissure, corpus callosum, fornix, and, anterior nucleus accumbens (MNI: 0.68) were selected. Using a hand-held dental drill, approximately 10 × 20 mm tissue blocks were dissected encompassing the corpus callosum as the dorsal landmark, anterior commissure as the ventral landmark, and fornix as the medial landmark. Tissue blocks were stored in sealed cryogenic bags at -80°C until cryosectioning.

### Tissue Processing for Anatomical Validation

Fresh frozen tissue blocks were placed inside the cryostat (Leica CM3050s) at -15°C for 30 minutes prior to cryosectioning. Tissue blocks were mounted on a round chuck with Optimal Temperature Compound (TissueTek Sakura, Cat #4583) for an additional 10-15 minutes. For each donor (*N*=4), a series of 30µm tissue sections were discarded until a flat surface was reached. The next 6-8 consecutive 10µm tissue sections were collected on microscope slides (VWR SuperFrost Microscope Slides, Cat #48311703). These slides were stored in slide boxes inside cryogenic bags at -80°C until ready for single molecule fluorescent *in situ* hybridization (smFISH) experiments.

### Single Molecule Fluorescent *in situ* Hybridization (smFISH) for Anatomical Validation and Co-Expression Visualization

RNAScope Multiplex Fluorescent Reagent Kit v2 (Advanced Cell Diagnostics [ACD], Cat #323100) and 4-Plex Ancillary Kit (ACD, Cat #323120) were used for smFISH. Tissue sections on microscope slides were fixed in 10% neutral buffered formalin solution (Sigma-Aldrich, Cat #HT501128) for 30 minutes at RT then underwent ethanol dehydration steps with 50%, 70%, and 100% ethanol solutions. Following dehydration, tissue sections underwent permeabilization treatments with hydrogen peroxide for 10 minutes at RT and protease IV for 30 minutes. Tissue sections on microscope slides were incubated for 2 hours at 40°C with commercially available probes. The slides were stored in 4x saline sodium citrate (Quality Biological, Cat #351–003-101) at 4°C for up to 72 hours. Probes were amplified and fluorescently labeled with fluorescent opal dye (Akoya Biosciences [AB]) diluted in TSA Buffer (ACD, Cat #322809) at a 1:500 ratio. For anatomical validation, the following probe and fluorescent opal dye combinations were used: Opal 520 (AB, Cat #FP1487001KT) for *MBP* (ACD, Cat #411051-C4), Opal 570 (AB, Cat #FP1488001KT) for *ELAVL2* (ACD, Cat #540301-C3), Opal 620 (AB, Cat #FP1495001KT) for *PPP1R1B* (ACD, Cat #477021), and Opal 690 (AB, Cat #FP1497001KT) for *TRPC4* (ACD, Cat #1226941-C2). For visualization of *OPRM1* expression in the LS, the following probe and fluorescent opal dye combinations were used: Opal 520 (AB, Cat #FP1487001KT) for *MBP* (ACD, Cat #411051), Opal 570 (AB, Cat #FP1488001KT) for *OPRM1* (ACD, Cat #410681-C4), Opal 620 (AB, Cat #FP1495001KT) for *GAD1* (ACD, Cat #404031-C3), and Opal 690 (AB, Cat #FP1497001KT) for *TRPC4* (ACD, Cat #1226941-C2). For visualization of *FREM2* expression in the LS, the following probe and fluorescent opal dye combinations were used: Opal 520 (AB, Cat #FP1487001KT) for *MBP* (ACD, Cat #411051-C4), Opal 570 (AB, Cat #FP1488001KT) for *FREM2* (ACD, Cat #482841), Opal 620 (AB, Cat #FP1495001KT) for *ELAVL2* (ACD, Cat #540301-C3), Opal 690 (AB, Cat #FP1497001KT) for *TRPC4* (ACD, Cat #1226941-C2). After a brief wash step, DAPI staining was applied on each slide for 20 seconds then coverslipped with microscope slide cover glass (Fisher Scientific, Cat #22-050-232) and 80uL Fluoromount-G (ThermoFisher, Cat #00-4958-02). Slides were allowed to dry at RT for 24 hours prior to imaging.

### Imaging for Anatomical Validation and Co-Expression Visualization

For anatomical validation, the imaging protocol from Ramnauth et al, 2023 [27] was implemented. Briefly, images were acquired using AX Nikon Ti2-E confocal fluorescence microscope equipped with NIS-Elements (v5.42.02) with 2x magnification (Nikon PLAN APO λ D 2x/0.1 objective) with a pinhole of 1.0 AU and the following laser power (LP) and gain (G) settings: DAPI: 27.00LP/18.00G, Opal 520: 28.29LP/25.00G, Opal 570: 25.00LP/20.00G, Opal 620: 25.00LP/20.00G, Opal 690: 25.00LP/25.00G, Lipofuscin: 28.29LP/25.00G. Images were uniformly gamma-adjusted (gamma = 0.5) for visualization purposes. For the *FREM2* and *TRPC4* coexpression validations, the following image acquisition settings were used: Nikon Plan Apo Lambda S 40x/0.9 objective with a pinhole of 1.0 AU, DAPI: 2.4LP/7.7G; Opal 520: 2.0LP/5.6G; Opal 570: 1.0LP/1.5G; Opal 620: 1.0LP/3.1G; Opal 690 4.8LP/4.0G; Lipo 2.0LP/19.4G. For the *OPRM1* presence validations, the following acquisition settings were used: Nikon Plan Apo Lambda S 40x/0.9 objective with a pinhole of 1.0 AU, DAPI: 2.4LP/5.6G; Opal 520: 1.8LP/2.7G; Opal 570: 1.3LP/1.0G; Opal 620: 1.3LP/3.5G; Opal 690 3.5LP/1.0G; Lipo 1.8LP/14.4G.

### snRNA-seq Data Generation

After anatomical validation, the 3 out of 4 fresh frozen tissue block samples containing the broadest extent of the LS were selected for snRNA-seq data generation. For each donor (*N*=3), 1-2 sections of 10µmtissue were discarded to reach a flat surface, then the adjacent 3 sections of 10µm tissue were collected on microscope slides (VWR SuperFrost Microscope Slides, Cat #48311703). Slides were stored in a slide box in -80°C. Prior to sample collection, each block was scored using a Single Edge Razor Blade (ULINE, Cat #H-595B) to minimize contamination from neighboring brain regions. The adjacent 3-5 100um tissue sections were collected in a prechilled 2mL microcentrifuge tube (Eppendorf Protein LoBind Tube, Cat #22431102) to total a volume of approximately 40 mg for each donor. Collected samples were stored in -80°C until ready for nuclei sorting. The next 3 sections of 10 micron tissue sections were collected on microscope slides then stored in a slide box in - 80°C until ready for validation experiments using smFISH.

Nuclei from each donors’ tissue samples were extracted as previously described [13]. Nuclei EZ Lysis Buffer (Millipore Sigma, Cat #NUC101) was chilled on wet ice then added to each 20mL PCR tube containing frozen tissue. Each tissue sample was homogenized using separate pairs of glass douncers and pestles per donor then filtered through 70µm mesh filter on separate 50 mL conical tubes. Filtrate was centrifuged at 500 rcf for 5 minutes at 4°C. Resulting pellets were washed with a total volume of 3 mL EZ Lysis Buffer. Final pellet was washed 3 times with wash buffer solution (1x PBS, 1% BSA, and 0.2U/uL RNAse Inhibitor). Nuclei underwent a 30 minute incubation on wet ice with Alexa Fluor 488-conjugated anti-NeuN-AF488 (Millipore Sigma, #MAB377X) diluted in dilution wash buffer at a 1:1000 ratio. Nuclei were labeled with propidium iodide (PI) diluted in a staining buffer solution (1x PBS, 3% BSA, and 0.2U/uL RNase inhibitor) at a 1:500 ratio. Samples were filtered through 35µm filter top FACS tubes then underwent fluorescent activated nuclear sorting (FANS) on a Bio-Rad S3e Cell Sorter. Gating criteria were set to select for whole, singlet nuclei by forward/side scatter then G0G1 nuclei by PI fluorescence and neuronal nuclei by Alexa Fluor 488 fluorescence. For each donor, approximately 6000 nuclei were collected, then the gating was adjusted to select for whole, singlet nuclei based on PI fluorescence for a 2:1 enrichment of neuronal nuclei. Nuclei were collected into 23.1uL master mix solution from 10x Genomics Single Cell 3’ Reagent kit (without reverse transcriptase). Reverse Transcription enzyme and water were added to full volume. cDNA and libraries were prepared according to the instructions provided by the manufacturer (10x Genomics Chromium Next GEM Single Cell 3’ Reagent Kits v3.1 Dual Index, CG000315 RevE) then sequenced on the Illumina NovaSeq6000 at the Johns Hopkins University Single Cell & Transcriptomics Core.

### snRNA-seq Raw Data Processing

FASTQ files for snRNA-seq sample libraries from 3 donors were aligned to the human genome (GRCh38/Hg38, Ensembl release 98), using 10x Genomics’ software [28], `cellranger count` (version 7.2.0). Raw feature-barcode files were analyzed in R v4.4.0, using the Bioconductor suite [24,25] of single-cell analytical packages version 3.19. Droplets containing nuclei were identified using the `emptyDrops()` function from the DropletUtils [29,30] package v1.24.0 using a data-driven `lower` threshold. Next, high quality nuclei were identified as those with less than 5% of reads mapping to the mitochondrial genome. Furthermore, following mitochondrial mapping rate thresholding, sample specific median absolute deviation (MAD) thresholds were identified for the number of UMI counts and detected genes per cell. The median value of the number of UMI counts and detected genes per cell for one particular sample (“Sample 1”) was much higher than the other two samples, and thus using an adaptive threshold for both UMI counts and detected features would have removed several hundred high quality nuclei. Therefore, only a UMI count threshold and mitochondrial mapping rate was used to remove low quality quality nuclei from Sample 1. In addition to thresholds for mitochondrial rate mapping, total UMI counts, and detected features, preliminary dimensionality reduction and clustering identified a cluster of low quality neuronal nuclei. Particularly, nuclei within this cluster exhibited expression of neurons, such as *GAD1, SYT1*, and *SNAP25*, but had abnormally low number of UMIs and detected genes. These nuclei were removed from further analysis. Following removal of low quality nuclei, sample specific doublet scores were calculated using the `computeDoubletDensity()` from the scDblFinder [31] v1.18.0 to identify any doublet driven clusters. These QC measures resulted in a final total of 9,225 nuclei with an average UMI count of ∼23,027 per nucleus and an average number of detected features per nucleus of ∼5,068.

### Feature Selection, Dimensionality Reduction, and Clustering

Feature selection was conducted by first calculating binomial pearson residuals. The top 2000 highly deviant genes (HDGs) were identified by ranking each gene according to its total deviance. Principal component analysis (PCA) was then performed within the 2000 HDG feature space using the binomial pearson residuals [32]. To avoid batch effects driven by donor/sample, we performed batch correction with mutual nearest neighbors within the reduced principal component space using the `reducedMNN()` function within the batchelor [33] package v1.20.0. Graph based clustering was then performed within the mutual nearest neighbors, PC-reduced space using `k=20` nearest neighbors and the Louvain method for community detection [34], yielding 26 preliminary clusters. Log2-normalized counts were calculated by first calculating size factors with the `computeSumFactors()` function provided by scran [35] v1.32.0, followed by log normalization with the `logNormCounts()` function provided by scuttle [36] 1.14.0. To visually inspect clusters, we generated a *t*-distributed Stochastic Neighbor Embedding (t-SNE) [37] using the top 50 mutual nearest neighbors-corrected PCs. Cluster identity was determined using previously identified transcriptional markers of cell types within the striatum [38–40], medial septum [14], and lateral septum [41]. Initial clustering identified 2 oligodendrocyte populations that were merged to generate a single oligodendrocyte cluster.

### Identification of Cluster Specific Marker Genes

Cluster specific marker genes were identified using an approach previously described with minor modifications. Briefly, log_2_-normalized counts were used to calculate two sets of statistics. The first set implemented the `findMarkers()` function provided by the scran [35] package v1.32.0 to perform pairwise *t*-tests and identify differences between each cluster. The second set utilized the `findMarkers_1vAll()` function provided by the DeconvoBuddies [42] package v0.99.0 to perform cluster-versus-all-other-nuclei testing.

### Comparison of Transcriptionally Distinct Cell Populations Across Human and Mice

To begin comparing transcriptomic profiles of transcriptionally distinct cell populations across humans and mice, we first conducted cluster-versus-all-other-nuclei testing using the `findMarkers_1vAll()` function provided by the DeconvoBuddies package v0.99. Shared homologs were then identified using JAX lab homology report (http://www.informatics.jax.org/downloads/reports/HOM_AllOrganism.rpt) and the “DB.Class.Key” identifier. For initial cross-species analysis focused on the correlation of every human and mouse cluster, we used the 917 shared homologs of the top 100 marker genes for each human or mouse cluster. All 16,574 shared homologs between human and mouse were used for the identification of conserved standardized log fold changes of shared homologs using the `cor()` function provided by base R. Human LS cell types were compared to the 10x genomics whole mouse brain taxonomy using MapMyCells [43], an interactive online tool provided by the NIH’s Brain Initiative Cell Census Network and the Allen Institute. First, human-mouse orthologs were identified using the conversion table provided by Allen Institute’s GeneOrthology package. Raw count matrices including all orthologs were uploaded to MapMyCells and the hierarchical mapping algorithm was used.

## RESULTS

### Characterization of transcriptionally distinct cell types within the human lateral septum

The LS is a bilateral structure situated around the midline of the brain ventral to the corpus callosum, extending between the fornix (medially) and the lateral ventricle (laterally). During the flash freezing of postmortem human brain tissue, the lateral ventricle collapses, shifting LS location to be adjacent to the caudate nucleus (CN) which is lateral to the lateral ventricle. In the human brain, the LS does not have clear anatomical boundaries between the medial septum (MS, medially), the bed nucleus of stria terminalis (BNST, laterally), and the ventral striatum (anterior-ventrally). These regions are transcriptionally and functionally distinct from the LS and vary along the anterior-posterior (A-P) axis. To ensure that dissections were enriched for LS rather than the neighboring regions, we performed RNAscope smFISH using regional markers identified in the mouse [39,41]: *TRPC4* to demarcate the LS, ELAV like RNA binding protein 2 (*ELAVL2*) to demarcate the MS, protein phosphatase 1 regulatory inhibitor subunit 1B (*PPP1R1B*) to demarcate the striatum, and myelin basic protein (*MBP*) to identify surrounding white matter (**Fig. S1**). Following anatomical validation, we captured single nuclei from the LS of 3 human brain donors using the 10x Genomics Chromium platform (**Fig. 1a**). Following quality control (see STAR Methods), 9,225 nuclei were divided into 25 transcriptionally distinct cell types (**Fig. 1b, Table S2, Fig. S2)**. These clusters included 19 neuronal cell types that exhibit high expression of pan-neuronal marker genes including synaptotagmin 1 (*SYT1*), synaptosome associated protein 25 (*SNAP25)*, and RNA binding fox-1 homolog 3 (*RBFOX3*/NeuN) (**Fig. 1c,d,k)**. We also identified 6 non-neuronal clusters including oligodendrocytes, marked by myelin associated oligodendrocyte basic protein (*MOBP*) expression (**Fig. 1i,k**) and astrocytes, marked by glial fibrillary acidic protein (*GFAP*) expression (**Fig. 1j,k**). The LS is predominantly inhibitory, containing GABAergic neurons [13,44] that are marked by glutamate decarboxylase 1 (*GAD1*) and glutamate decarboxylase 2 (*GAD2*), genes encoding enzymes critical for the synthesis of gamma-aminobutyric acid (GABA) (**Fig. 1e,f,k**). Several distinct clusters exhibited expression of both solute carrier family 17 member 6 (*SLC17A6*/VGLUT2) and solute carrier family 17 member 7 (*SLC17A7*/VGLUT1) (**Fig. 1g,h,k**). The MS contains glutamatergic neurons [45,46] suggesting that these cells may not have originated from the LS, but rather a neighboring brain region.

**Figure 1.**
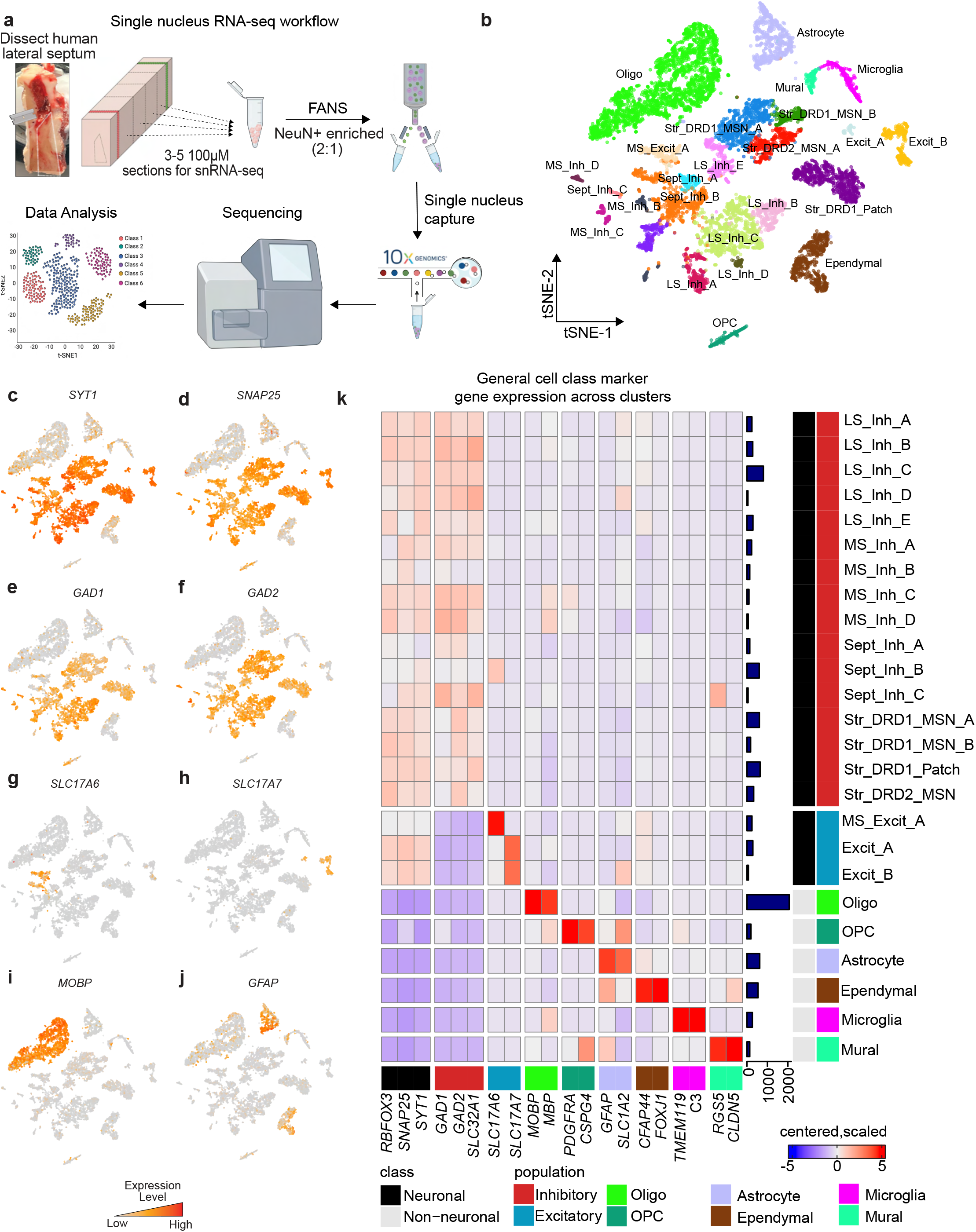
snRNA-seq of the human lateral septum (LS) identifies 25 transcriptionally distinct cell populations. **a**, Description of workflow to dissect human LS, dissociate nuclei and perform droplet based single nucleus capture. **b**, t-distributed Stochastic Neighbor Embedding of 9,225 nuclei from human LS colored by cluster identity. Features plots demonstrating distribution of expression across cell types for **c**, *SYT1*, **d**, *SNAP25*, **e**, *GAD1*, **f**, *GAD2*, **g**, *SLC17A6*, **h**, *SLC17A7*, **i**, *MOBP*, and **j**, *GFAP*. **k**, Heatmap of expression values for general cell class marker genes used to determine cluster identity. Color of square corresponds to centered and scaled expression values (logcounts).

To determine the brain region of origin for each cluster, we used established marker genes (**Fig. 2, Fig. S3**). First, clusters representing septal nuclei were identified by expression of FXYD domain containing ion transport regulator (*FXYD6*; **Fig. 2a,b,e**), a gene that is abundantly expressed in mouse in both MS and LS [14]. Although this gene is detected in all neuronal clusters, it is most highly expressed in nuclei derived from the MS and LS (**Fig. 2a,b,e**). To delineate between the LS and MS, we queried expression of *TRPC4*, diacylglycerol kinase gamma (*DGKG*), and corticotropin releasing hormone receptor 2 (*CRHR2*), LS marker genes that were identified in rodent studies (**Fig. 2a,c,f**) [8,13,14,47]. Five clusters exhibited relatively high expression of these genes (LS_Inh_A, LS_Inh_B, LS_Inh_C, LS_Inh_D, LS_Inh_E) and were classified as LS neuronal subtypes. MS neuronal subtypes (MS_Inh_A, MS_Inh_B, MS_Inh_C, MS_Inh_D, MS_Excit_A) were identified using *ELAVL2*, adenosine deaminase RNA specific B2 *(ADARB2)*, and SRY-box transcription factor 6

**Figure 2.**
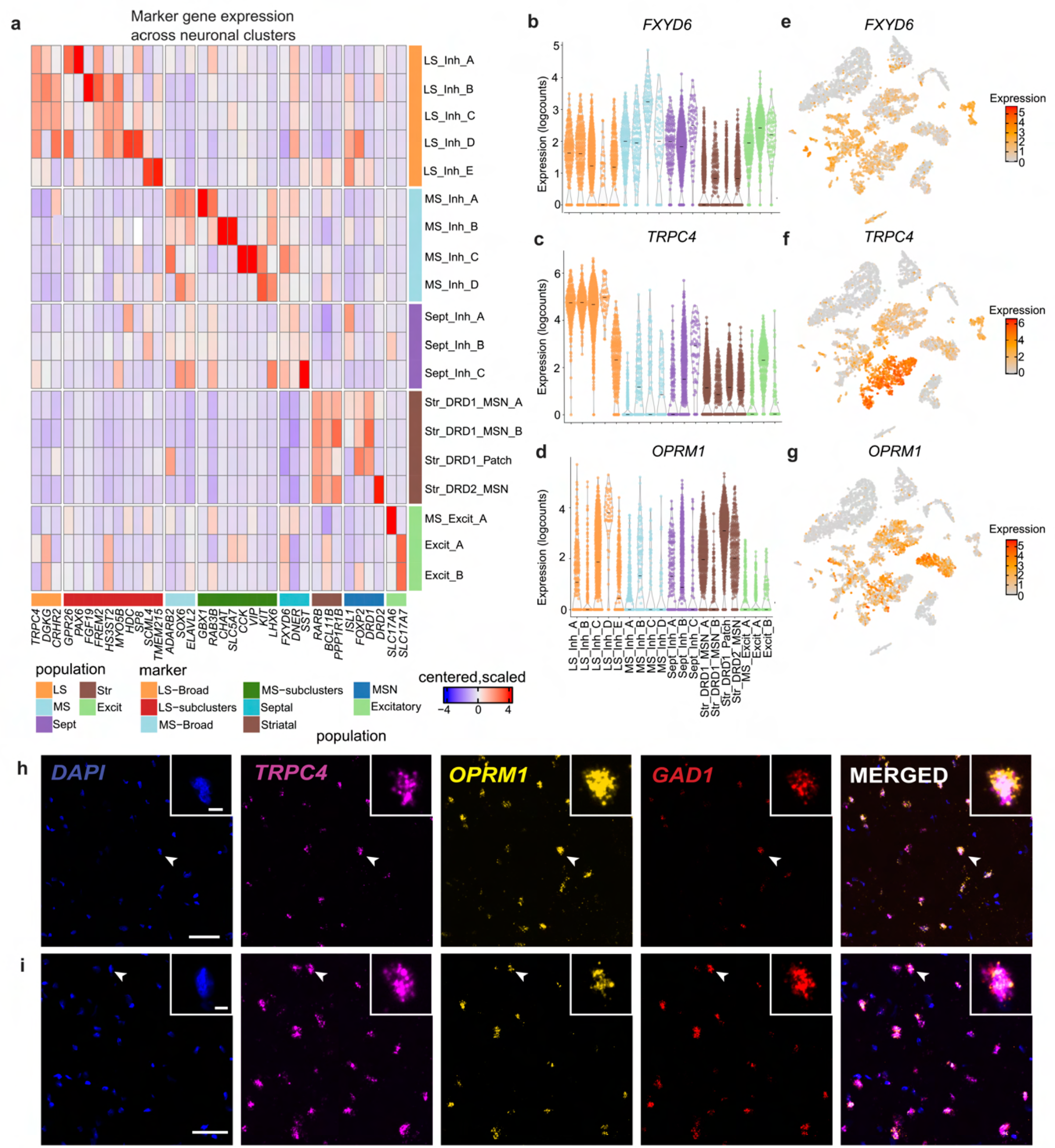
Identification of novel marker genes of human lateral septum (LS). a, Heatmap of expression values for neuronal clusters within the human LS, medial septum (MS), and striatum (Sir). Violin plots of expression values for **b**, *FXYD6*, **c**, *TRPC4*, and **d**, *OPRM1*. Feature plots demonstrating distribu tion of expression values fore, *FXYD6*, **f**, *TRPC4*, and **g**, *OPRM1*. **h**, Representative smFISH images demonstrating the coexpression of *TRPC4* and *OPRM1* transcripts in tissue section collected from the anterior portion **i**, posterior portion of LS from an independent donor. White bar indicates 50µm. White arrowheads indicate cells in inset.

*SOX6* (**Fig. 2a**). MS_Excit_A exhibits high expression of *SLC17A6* (**Fig. 1g,k, Fig. 2a**). *SLC17A6* marks a discrete population of excitatory MS neurons that is important for arousal and wakefulness in mouse models [48]. We also used a panel of molecular markers (retinoic acid receptor beta (*RARB*), BCL11 transcription factor B (*BCL11B*), and protein phosphatase 1 regulatory inhibitor subunit 1B *(PPP1R1B*)) to identify nuclei originating from the CN, or striatum (**Fig. 2b, Fig. S3**). Additionally, striatal neurons are both spatially and transcriptionally distinct [38–40,49–51]. Our dataset contains *DRD1*-expressing medium spiny neurons (MSNs) as denoted by expression of opioid receptor mu 1 (*OPRM1*), crystallin mu (*CRYM*), and semaphorin 5B (*SEMA5B*; **Fig. 2a, Fig. S3**).

### Identification of transcriptionally distinct neuronal subtypes within the human lateral septum

We next investigated LS neuronal subtype heterogeneity by identifying genes that are preferentially enriched within each subtype using a cluster-versus-all-other-clusters approach (**Fig. 2a, Table S3**). While this analysis will identify genes that are most abundant within each cluster, it will not specifically identify selective markers. For example, *FREM2* is highly expressed within the LS_Inh_B cluster (**Fig. 2a**), and is identified as a marker gene for this cluster. However, this gene is widely expressed in most LS clusters originating from the anterior and middle LS (LS_Inh_A, LS_Inh_B, LS_Inh_C, LS_Inh_D), and may represent an additional marker of the human LS (**Fig. 2a**). Similarly, Myosin VB (*MYO5B*) is preferentially enriched within LS_Inh_C, but is also expressed within LS_Inh_A, LS_Inh_B, LS_Inh_C, and LS_Inh_D (**Fig. 2a)**, and may represent an additional marker of the human LS. G protein-coupled receptor 26 (*GPR26*), which encodes an orphan GPCR that is important for satiety and feeding behaviors [52], and paired box 6 (*PAX6*) are both preferentially enriched within LS_Inh_A (**Fig. 2a)**. Myosin VB (*MYO5B*) is preferentially enriched within LS_Inh_B, while Sp8 transcription factor (*SP8*) is enriched within LS_Inh_D (**Fig. 2a**). Finally, Transmembrane protein 215 (*TMEM215*) and Scm polycomb group protein like 4 (*SCML4*) were identified as marker genes for LS_Inh_E. Cells within the LS_Inh_E cluster do not express canonical LS marker genes to the same extent as other LS clusters. However, cells within this cluster are not significantly enriched for any MS or CN markers (**Fig. 2a, Fig. S3**). Enrichment of LS marker genes and depletion of striatal and MS marker genes led us to classify LS_Inh_E as an LS neuronal subtype.

The µ-opioid receptor is expressed in several transcriptionally distinct neuronal subtypes in rodent LS [16], and regulates opioid-dependent behaviors that model specific aspects of human drug addiction [53,54]. We observe distributed expression of *OPRM1* across LS neuronal subtypes with preferential enrichment in the LS_Inh_D cluster (**Fig. 2d,g**). We validated *OPRM1* expression in the human LS from an independent brain donor using smFISH. We multiplexed probes for *TRPC4, GAD1*, and *OPRM1* in tissue sections originating from the anterior and posterior portions of the LS, and found co-expression of *TRPC4, GAD1*, and *OPRM1* (**Fig. 2h,i**). *OPRM1*-expressing neurons do not appear to be spatially organized as *OPRM1* punctae were identified within both anterior and posterior tissue sections. This experiment confirmed *OPRM1* expression in human LS neurons and validates the findings from the snRNA-seq analysis.

### Transcriptional signatures of discrete LS neuronal populations are conserved across mice and humans

We next investigated conservation of transcriptional signatures of LS cell types across human and mouse using two independent datasets. First, we utilized MapMyCells [43], an online application that allows users to compare their data to high quality, large-scale single cell sequencing datasets. Using the 10x Genomics whole mouse brain taxonomy dataset [43], we calculated the percentage of human cells from each cluster that map onto each broad mouse cell class. Almost 100% of each non-neuronal population map onto the same non-neuronal population in mouse (**Fig. S4a, Table S4**). A similar trend was observed with neurons originating from the LS_Inh_A, LS_Inh_B, LS_Inh_C, and LS_Inh_D populations mapping to GABAergic LS mouse neurons (**Fig. S4**). LS_Inh_E, a population of human LS neurons marked by the expression of *TRPC4* and the absence of many broad striatal markers, mapped to GABAergic populations in the mouse striatum (**Fig. S4**). We were particularly interested in identifying LS cell types and marker genes that are similar between human and mouse. Using a recently published transcriptional atlas of the mouse LS [13], we performed 1-versus-all differential expression testing for each gene per cluster within each dataset and identified shared homologs for the top marker genes of each cluster. Then, we calculated the pearson’s correlation coefficient (*r*) by correlating each gene’s standardized log-fold change within each cluster across species. Transcriptional signatures for non-neuronal populations such as microglia, oligodendrocytes, and ependymal cells are highly correlated between human and mouse (**Fig. 3a**). LS_Inh_E exhibited positive correlations with both mouse LS and striatal clusters (**Fig. 3a**). The Excit_A and Excit_B neuronal clusters are also highly correlated with mouse clusters originating from the tenia tecta, indusium griseum, and septohippocampal nucleus (**Fig. 3a**).

**Figure 3.**
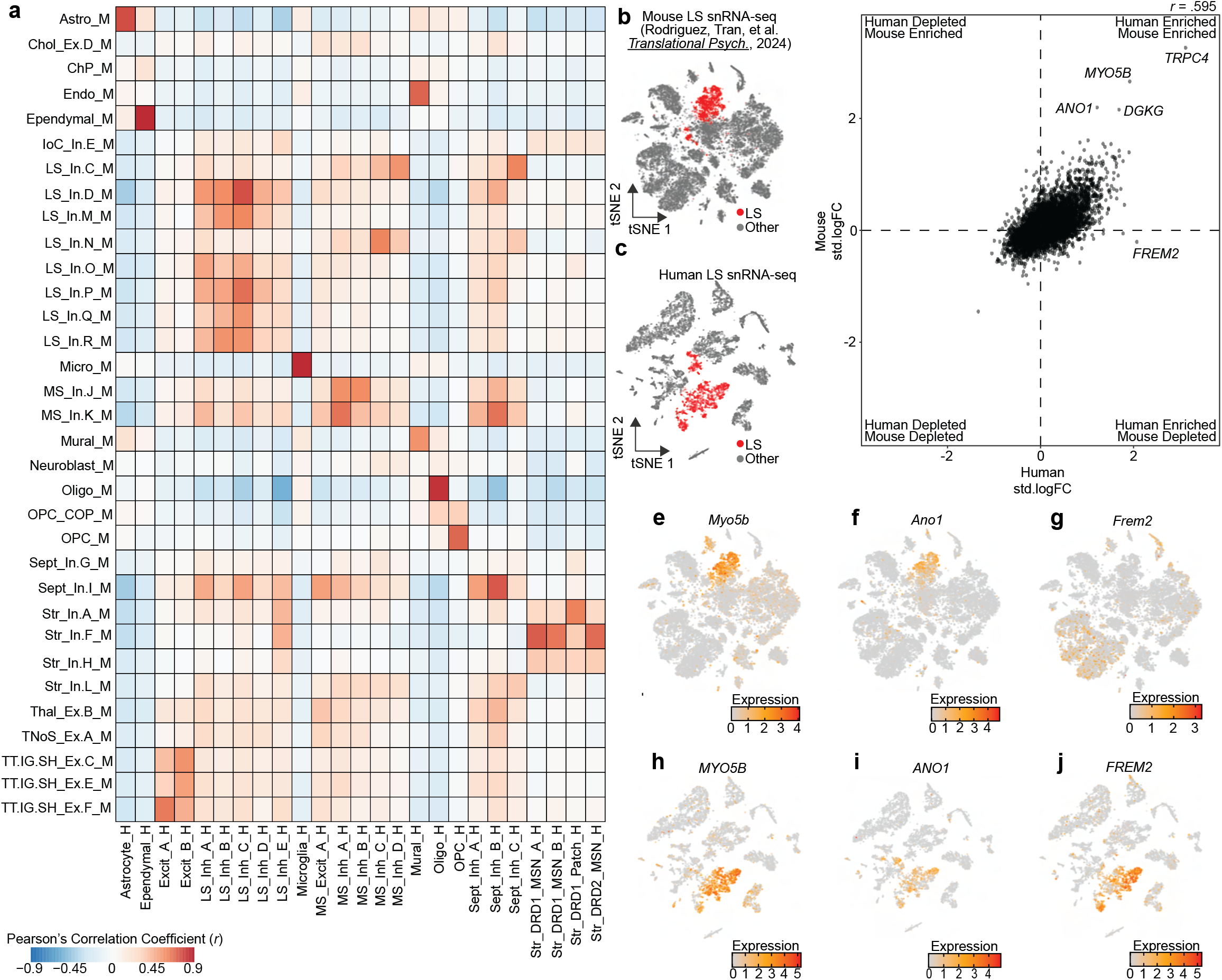
Identification of conserved transcriptional markers for lateral septum (LS) neurons. **a**, Heatmap of Pearson’s correlation coefficients calculated by correlating the standardized log fold changes of the top 100 marker genes from each cluster that has a homologous partner in each species. **b-c**, t-SNE of mouse and human snRNA-seq datasets containing LS neurons. Cells are colored by LS neuronal identity. **d**, Standardized log fold changes were calculated for all genes in LS neurons in mouse or human and correlated to identify conserved transcriptional markers. **e-g**, Feature plots demonstrating distribution of expression of *Myo5b, Ano1*, and *Frem2* in mouse. **h-j**, Feature plots demonstrating distribution of expression of *MYO5B, ANO1*, and *FREM2* in human snRNA-seq dataset containing the LS.

Interestingly, no particular LS neuronal cluster exhibited a high degree of conservation across species. Instead, LS neuronal clusters exhibited broad conservation (**Fig. 3a**), suggesting that primary markers of LS neurons are conserved while distinct marker genes for discrete subtypes may exhibit divergence across species. To identify both conserved and divergent marker genes for LS neurons across the human and mouse, we first collapsed all neuronal LS clusters within each species (**Fig. 3b,c**), and then recalculated the standardized log-fold change for all 16,574 shared homologs between human and mouse. Overall, gene enrichment within mouse and human neuronal LS clusters are correlated (*r* = 0.595, *p* < 2.2×10^−16^, **Fig. 3d**). As expected, *TRPC4* and *DGKG* were enriched in neuronal LS clusters in both the human and mouse (**Fig. 3d**). We also identified *ANO1* and *MYO5B* as strong markers of the LS in both humans and mice (**Fig. 3d-i**). Additionally, this analysis identified that *FREM2*, a marker gene for LS_Inh_B, was specifically enriched only within neuronal LS clusters from the human, and represents a potentially species-divergent LS marker gene (**Fig. 3g-j**).

### *In situ* validation of *FREM2* as a marker gene for human LS neurons

Identifying robust human LS markers is important as there are no clear anatomical landmarks that distinguish the LS from surrounding brain structures such as the MS (medially), NAc (ventrally), and the BNST (laterally). Thus, anatomical validation of tissue sections containing the LS relies on targeted amplification of biomolecules, such as mRNA transcripts, to ensure accurate dissection. Our cross-species analysis identified *FREM2* as a selective marker gene for human LS neurons (**Fig. 3d,j**). We validated this finding *in situ* by probing for *FREM2* in an independent brain donor (**Fig. 4a,d**). Because gene expression is spatially organized within the LS [47], we performed smFISH with probes for *TRPC4* and *FREM2* on one anterior and one posterior section and used myelin basic protein (*MBP*) staining to identify surrounding white matter regions for anatomical orientation (**Fig. 4b,e**). We found that *FREM2* is highly expressed in both the anterior and the posterior section, confirming it as a transcriptional marker of the LS (**Fig. 4b-f**). A high percentage of *TRPC4+* neurons are also *FREM2+*, further confirming this gene as a marker of human LS (**Fig. 4c,f)**. Interestingly, *FREM2* expression was comparatively lower than *TRPC4* and also exhibited differences across snRNA-seq clusters assigned to the LS (**Fig. 2a**). This difference in enrichment is further demonstrated by the standardized log-fold change as the fold change is higher for *TRPC4* than *FREM2* (**Fig. 3d**).

**Figure 4.**
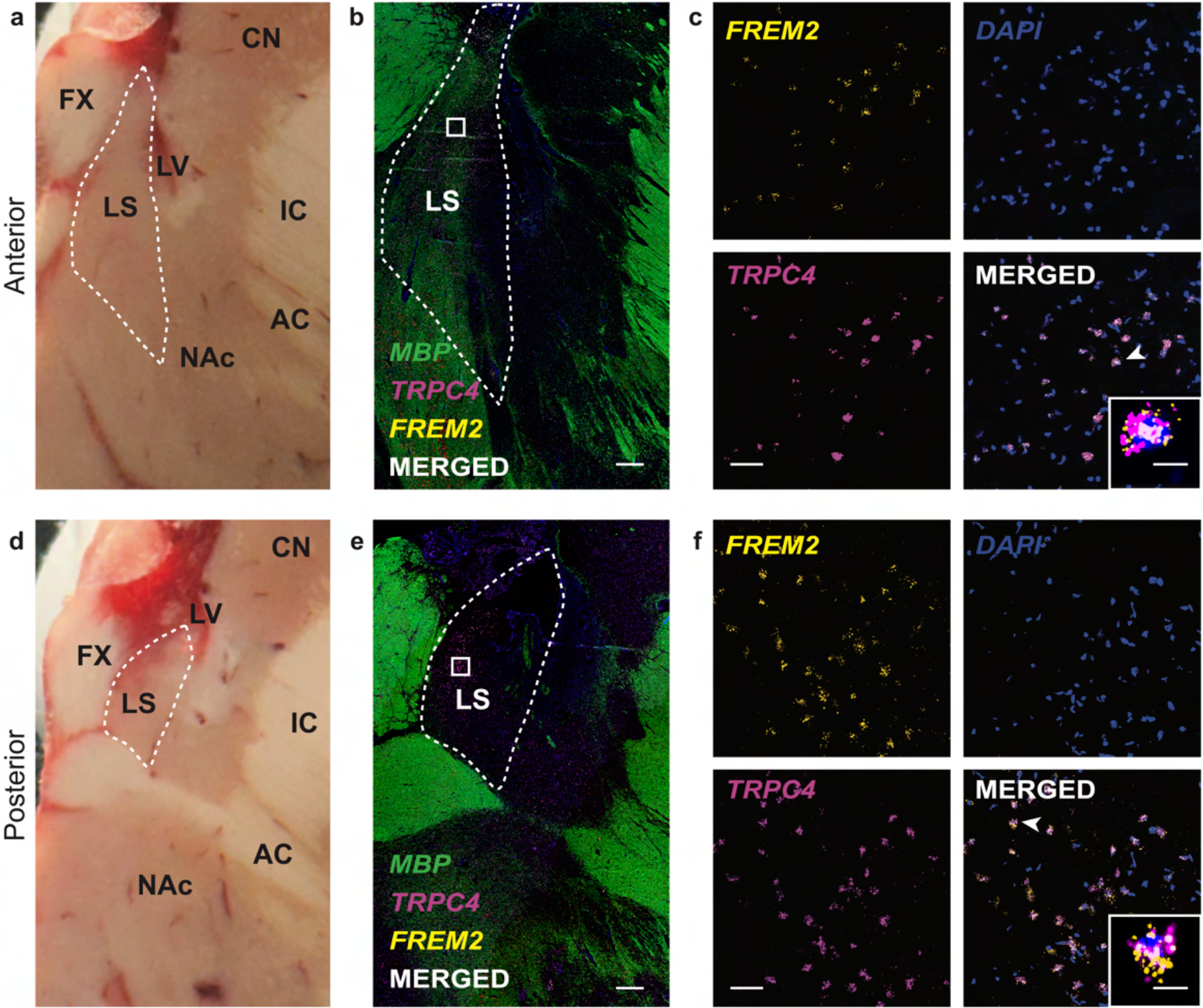
*In situ* validation of *FREM2* expression in the lateral septum (LS). **a, d**, Postmortem human brain block from a donor not included in the snRNA-seq study shown at the level of anterior LS **(a)** and posterior LS **(d)**. LS is outlined by the dashed white line. CN - caudate nucleus; FX-fornix; LS - lateral septum; IC - internal capsule; AC - anterior commissure; NAc - nucleus accumbens. **b, e**, 2x magnification smFISH images at the anterior LS **(b)** and posterior LS **(e)** illustrating expression of *TRPC4* and *FREM2. MBP* expression is included for anatomical visualization of white matter. LS is outlined by the dashed white line. White squares indicate the approximate location of the corresponding 40x images. White bar indicates 1000µm. **c, f**, 40x magnification smFISH images of the area demarcated by white squares in 2x images from anterior LS **(c)** and posterior LS **(f)**. White bar indicates 50µm. Merged image shows *TRPC4* and *FREM2* co-expressing cells in white. Inset depicts a zoomed in image of a single neuron indicated by the white arrowhead. White scale bar of the inset indicates 10µm.

## DISCUSSION

The LS regulates evolutionarily conserved behaviors that are frequently dysregulated in neuropsychiatric and neurodevelopmental disorders. Despite its essential function in regulating social and affective behaviors, the molecular and transcriptional heterogeneity of cell types within the LS has only recently been addressed. snRNA-seq studies of the mouse LS identified dozens of transcriptionally distinct neuronal subtypes that are implicated in social and affective behaviors [13–16]. For example, our recent snRNA-seq study of the mouse LS demonstrated that transcriptionally distinct neuronal subtypes are differentially regulated by TrkB signaling [13], providing insight about cell type-specific function of brain-derived neurotrophic factor (BDNF)-TrkB in the LS in social recognition. Additionally, multimodal studies identified electrophysiological properties specific to transcriptionally distinct neuronal subtypes across the rostral-caudal extent of the mouse LS [15]. While molecular and cellular properties of LS neurons can be functionally interrogated in rodent models, a major limitation is the ability to relate these findings to analogous populations of neurons within the human brain.

To address this limitation, we used snRNA-seq to generate the first comprehensive molecular profiles of human LS cell types. Our study identified 19 transcriptionally distinct neuronal subtypes, including 5 that are LS specific (**Fig. 1b**). The human LS is surrounded by two GABAergic brain regions that exhibit high levels of expression of *GAD1* and *SLC32A1*, the gene encoding the vesicular GABA transporter. Thus, accurate identification of human LS populations cannot rely on the presence of GABAergic markers alone. Hence for our dissections, we primarily relied on expression of *TRPC4*, a gene that was identified as a marker of the mouse LS [13]. smFISH studies confirmed that human LS neurons exhibited high expression of *TRPC4* (**Fig. S1**). The snRNA-seq data also confirmed the presence of this gene within LS (**Fig. 2a,c,f**). In addition to LS-specific marker genes, we also found that transcriptional markers of the broad septal complex in the mouse, *FXYD6* and *DGKG*, were enriched within neuronal clusters originating from human LS and MS (**Fig. 2a,b,e**). Interestingly, *DGKG* was specifically enriched within human LS neuronal clusters (**Fig. 2a**). The enrichment of mouse LS marker genes in human LS populations suggest that region-specific transcriptional markers may be conserved across species.

To investigate cross-species enrichment of marker genes, we first compared the standardized log fold change for each gene on a per cluster basis for human and mouse clusters. Correlation of standardized log fold changes for homologous genes between clusters demonstrated that human and mouse LS neuronal clusters are broadly conserved, without any specific human cluster mapping to one mouse cluster (**Fig. 3a**). This is in contrast to glial populations which exhibit high correlation coefficients across the two species (**Fig. 3a**). Broad conservation suggests that marker genes of the entire LS, instead of the markers of discrete neuronal subtypes, are conserved across species. We next correlated LS-specific standardized log fold changes between human and mouse. This analysis found that *TRPC4* was highly conserved, confirming the findings from smFISH studies in both human (**Fig. S1**) and mouse [13]. *FREM2* was identified as a human-specific LS marker gene and validated via smFISH results (**Fig. 3d,j, Fig. 4a-c**). To our knowledge, no other studies have identified *FREM2* as a marker gene of the LS. While this analysis represents the first cross-species study of transcriptomic signatures of LS neurons, additional integration with recently generated whole brain atlases is needed. The current dataset includes several tissue sections from various points along the anterior-posterior axis that are combined into a single sample. Spatial information is required to accurately integrate with mouse datasets in which the transcriptomic differences along the anterior-posterior axis are better understood. This will require additional experiments in which spatial information is retained within snRNA-seq experiments or using spatially-resolved transcriptomic technologies.

Our dataset includes 5 transcriptionally distinct LS neuronal subtypes (LS_Inh_A, LS_Inh_B, LS_Inh_C, LS_Inh_D, LS_Inh_E). All neuronal subtypes exhibit expression of *TRPC4* and *DGKG*, but are also enriched for cluster-specific genes (**Fig. 2a**). *OPRM1*, the gene encoding the µ-opioid receptor, was expressed in multiple LS clusters, but primarily enriched within the LS_Inh_D cluster (**Fig. 2a,d**). This finding is in agreement with recent snRNA-seq of the mouse LS that found *Oprm1* to be expressed in several LS neuronal subtypes [16]. Opioid-related deaths continue to climb, thus identifying the neuronal subtypes that may regulate opioid addiction-related behaviors is critical for understanding how drugs impact the brain and the development of future therapeutics. Corticotropin releasing hormone (CRH) signaling within the LS is critically important for social behaviors in mice [8]. Interestingly, LS_Inh_A, LS_Inh_B, LS_Inh_C, and LS_Inh_D are all enriched for *CRHR2* (**Fig. 2a & Fig. S3**), a gene encoding a receptor for CRH signaling. LS_Inh_E was enriched for *TRPC4*, but also exhibited expression of transcriptional markers of striatal MSNs *BCL11B, PPP1R1B*, and *ISL1* [39,40,51] (**Fig. 2a**). A large portion of the neurons in LS_Inh_E originates from a single donor in which the tissue punches were taken from a more posterior portion of the LS, suggesting that this cluster may represent a cell type that marks the transition from LS to striatum (**Table S2, Fig. S1**). The presence of a transcriptionally distinct LS neuronal subtype originating from a specific position along the rostral-caudal axis suggests that some clusters may be both transcriptionally and spatially distinct, similar to observations in mouse. Additionally, LS_Inh_A is significantly enriched for *PAX6*, a gene that marks the ventral LS within the mouse brain [55]. Thus, LS_Inh_A may represent a transcriptionally and spatially distinct population of LS neurons. It is important to note that this dataset does not contain neurotensin neurons. Within the mouse, neurotensin neurons regulate social approach behavior [12] and are modulated by opioid withdrawal [16]. However, the absence of neurotensin neurons may be due to technical issues with capture of *NTS* mRNA. A recently published snRNA-seq study of the mouse LS from our group utilized the same 3’ mRNA capture technology used for this study and likewise failed to identify neurotensin neurons [13]. Thus, identification of human neurotensin neurons may require additional studies focused on the amplification of *NTS* mRNA via smFISH or spatial transcriptomic approaches.

In conclusion, single nucleus resolution allowed us to identify cluster-specific marker genes, such as *OPRM1*, while also providing the ability to identify marker genes for the broad LS. While snRNA-seq provides unprecedented resolution into the transcriptome of LS neurons, it does not provide information regarding its spatial organization. Mouse studies have identified that the LS is composed of non-overlapping spatial domains that contain transcriptionally distinct LS nuclei [15,47]. Our data suggests that the human LS is also composed of transcriptionally and spatially distinct neuronal cell types. Future studies of the human LS will require spatial transcriptomics to understand how transcriptionally distinct cell types are spatially organized, both within a single tissue section and along the A-P axis.

## DATA AVAILABILITY

Raw sequencing data is publicly available at https://www.ncbi.nlm.nih.gov/bioproject/PRJNA1093410. The R object containing count matrices and cell identities is available for download at https://research.libd.org/globus/ under `jhpce#ls_molecular-profiling_sce`. All code for analyzing the data is publicly available at https://github.com/LieberInstitute/ls_molecular-profiling. Any additional information required is available from the lead contact upon request.

## ACKNOWLEDGEMENTS

A portion of Figure 1 was created with BioRender.com. We thank the LIBD neuropathology team, particularly James Tooke and Amy Deep-Soboslay, for curation of the brain samples and assistance with tissue dissections, and the families of the brain donors for their generosity. We thank Kristen Maynard for feedback on the manuscript. Finally, we thank the families of Connie and Steve Lieber and Milton and Tamar Maltz for their generous support.

## AUTHOR CONTRIBUTIONS

Conceptualization: KM, SCP

Data Curation: RAP, RAM

Formal Analysis: RAP

Investigation: SO, SVB, YD, SCP

Validation: SO, SVB, YD

Resources: JEK, TMH

Software: RAM

Visualization: RAP, SO, RAM, SVB

Project Administration: KM, SCP

Supervision: KM, SCP, SCH

Funding acquisition: KM

Writing – original draft: RAP, SO, SVB

Writing – review & editing: RAP, SO, SVB, SCH, SCP, KM

## Supplementary Tables

**Supplementary Table 1. Donor demographic information**. Demographic information on brain donors including Brain ID, diagnosis information, sex, race, age, and manner of death. An additional column containing the type of experiment (RNAscope vs snRNA-seq) the tissue was used for is also included.

**Supplementary Table 2. Sample representation by cluster**. snRNA-seq data was generated from three different donors and identified 25 transcriptionally distinct cell types. This table contains the number of nuclei within each cluster by the sample/donor.

**Supplementary Table 3. Differential Expression Statistics**. Differential expression testing was performed via 1-versus-all or pairwise testing on a per-cluster basis. Additionally, 1-versus-all testing was performed between LS nuclei versus all other nuclei. This table contains three sheets containing differential expression statistics for genes with an adjusted *p*-value < 0.05.

**Supplementary Table 4. MapMyCells Output Statistics**. MapMyCells was used to identify cell types within the mouse brain that were similar to cells dissected and sequenced from the post-mortem human brain. This table contains output statistics from the MapMyCells analysis including bootstrap probability.

