## Supplementary figures and images for "Transcriptomic characterization of human lateral septum neurons reveals conserved and divergent marker genes across species"

### Supplemental Figure 1

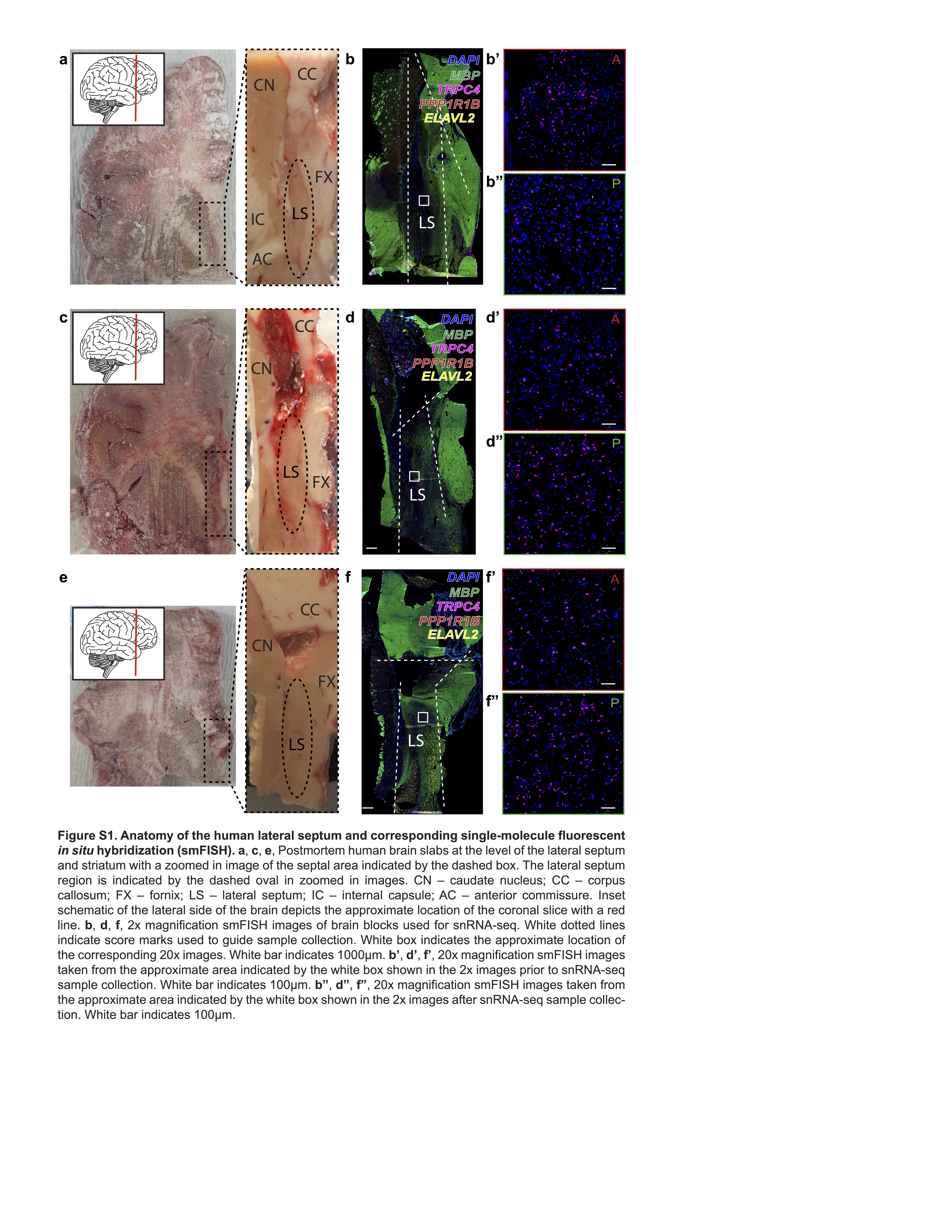
