## Supplemental Figure 2 for "Transcriptomic characterization of human lateral septum neurons reveals conserved and divergent marker genes across species"

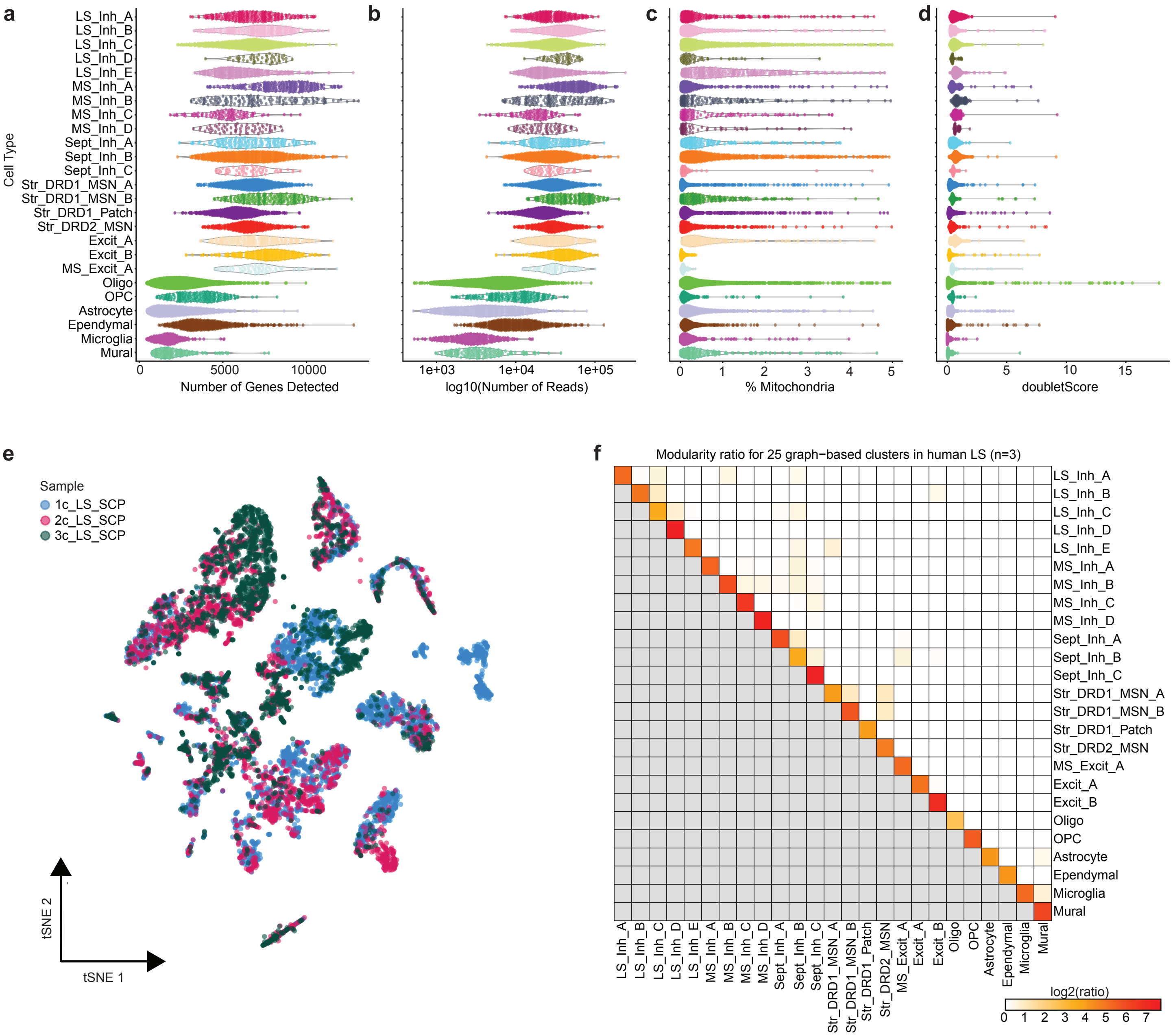


**Figure S2**. Cluster and nuclei quality control. Violin plots of various quality control metrics such as **a**, number of genes detected **b**, number of reads **c**, percent of reads mapping to the mitochondrial genome and **d**, doublet score. **e,** t-SNE with nuclei colored by donor/sample. **f**, Heatmap of pairwise modularity scores for 25 clusters from human LS single nucleus RNA-sequencing.
