## Supplemental Figure 3 for "Transcriptomic characterization of human lateral septum neurons reveals conserved and divergent marker genes across species"

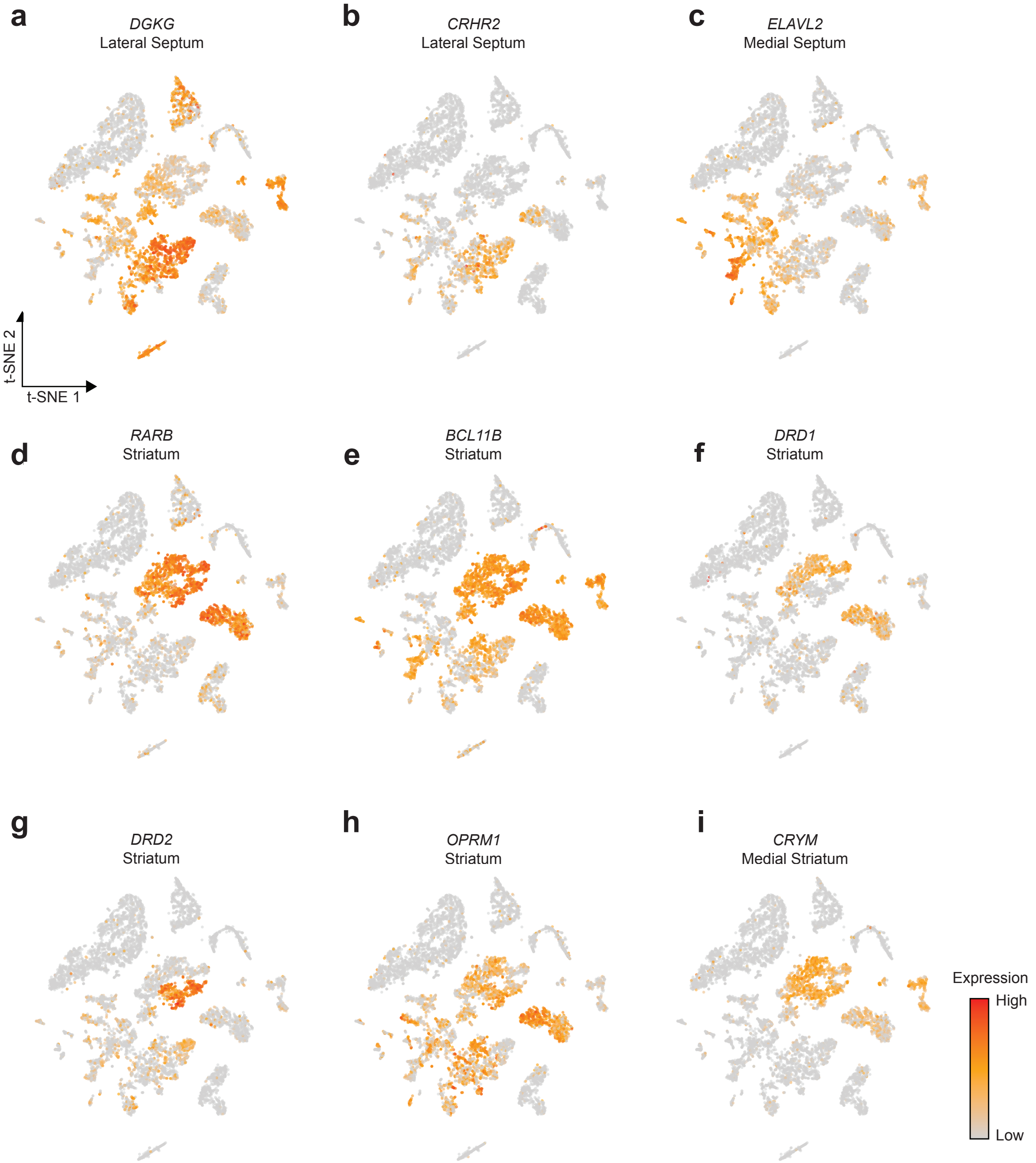


**Figure S3**. Distribution of expression of septal and striatal marker genes. Feature plots for **a**, *DGKG*, **b**, *CRHR2*, **c**, *ELAVL2*, **d**, *RARB*, **e**, *BCL11B*, **f**, *DRD1* **g**, *DRD2*, **h**, *OPRM1*, **i**, *CRYM*. Titles include gene name and brain region where gene is most highly expressed.
