## Supplemental Figure 4 for "Transcriptomic characterization of human lateral septum neurons reveals conserved and divergent marker genes across species"

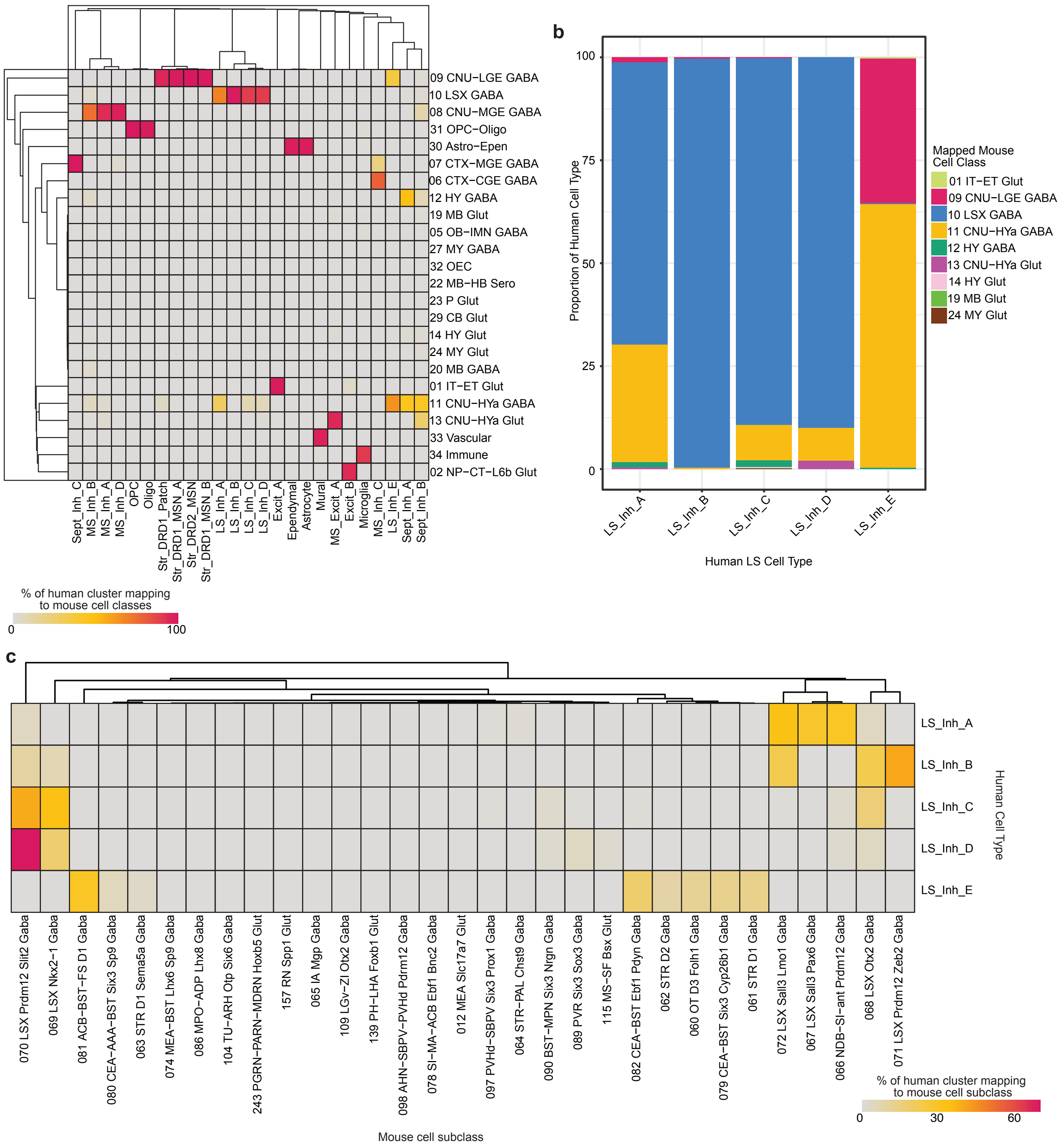


**Figure S4**. Cross-species analysis with MapMyCells. **a**, Heatmap of the percentage of each human cell cluster mapping to broad mouse cell classes. **b**, Barplot for the proportion of each human LS neuronal cell type mapping to broad mouse cell classes. **c**, Heatmap of the percentage of each human LS neuronal cell type mapping to mouse subclasses.
